# *In vivo* Stability and Biodistribution of Liposome Coated with SlpB from *Levilactobacillus brevis*

**DOI:** 10.1101/2023.04.06.533723

**Authors:** Zheng Lin Tan, Naoyuki Yamamoto

## Abstract

SlpB from *Levilactobacillus brevis* offers a solution to stabilise liposome in gastrointestinal tract, and to target intestinal APCs in Peyer’s patches, rendering it a powerful tool for oral delivery of drugs, and to yield the benefits provided by oral delivery. However, the stability of SlpB-coated liposome (SlpB-LP) and its distribution in tissues were not characterized. In this study, we have demonstrated that SlpB-coating could improve the stability of liposome in gastrointestinal tract, and facilitate specific uptake of liposome into Peyer’s patches, but not intestinal, nor intestinal mucosa. Furthermore, we have shown that uptake of SlpB-LP into Peyer’s patches enhanced bioavailability of drugs, which have resulted in 427.65-fold increase in bioavailability and at least 2.41-fold decrease in retention of fluorophore in liver where drug metabolism takes places, to a degree which approximate control group. In conclusion, this study shows that SlpB could increase stability of liposome in gastrointestinal tract, increase specific uptake of liposome into Peyer’s patches, and improve bioavailability.

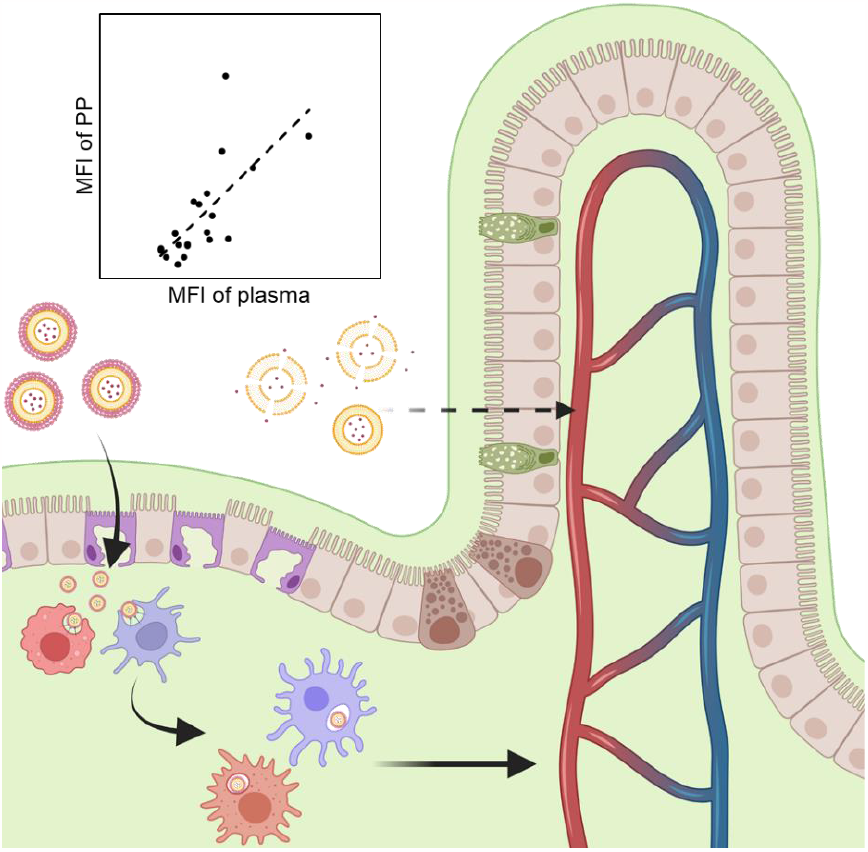

## Introduction

Oral delivery of drugs is regarded as a more preferential route of drugs administration due to high patients’ compliance. For immunomodulators, oral delivery allow access to the largest immune organ in mammal, i.e., Peyer’s patches which house 70% of immunocyte [1,2], and allow absorption of drugs into lymph, which can bypass first pass elimination [3]. However, oral delivery of drugs is complicated due to mechanical stress, diverse pH, salinity, poor aqueous solubility, short residence time, and existence of mucosal barrier in gastrointestinal tract [4].

Various formulation of have been developed to enable oral delivery of drugs which includes polymeric nanoparticles [5,6], clay [7], and bacteria [8,9] by exploiting the capability of microfold (M) cells in Peyer’s patches to transcytose microparticles [10]. However, uptake of solid carriers into Peyer’s patches is slow, due to the larger size and higher rigidity of solid nanoparticles.

Liposome is a fascinating drug vehicle for its capability to load both hydrophobic and hydrophilic drugs, and higher uptake efficiency compared to other carriers due to difference in uptake mechanism, i.e., via lipid endocytosis. Although some formulation, e.g., Eudragit® L-100 which is a pH sensitive polymer dissolute at pH > 6.0 could protect liposome against degradation in stomach, liposome is subject to emulsify by bile solution and hydrolyse by lipase contained in pancreatic juice. Furthermore, in practical application, liposomes are not fed directly into stomach. Mouth and oesophagus which have pH > 6.0 provide condition for dissolution of protective layer. Slps from probiotic lactobacilli are attractive candidate to protect liposome against gut environment [11] and to target antigen presenting cells (APCs) due to their capability to interact with immunocytes [12,13]. Prior studies have shown that Slp from *L. helveticus* could protect liposomes against degradation and bind to mucosal layer of stomach and intestine. This property of Slp-coated liposome (Slp-LP) allow effective delivery of vaccine via oral administration. However, delivery of liposome into Peyer’s patches was not reported, probably due to dissolution of liposome in mucosal layer through Slp-mucosa interaction. By replacing Slp from *L. helveticus* (*L. acidophilus* group lactobacilli) with SlpB from *Lv. brevis*, we have demonstrated that SlpB-LP could facilitate and enhance delivery of liposome to APCs in Peyer’s patches [14], which provides a fascinating tool for gut immunocytes-targeting drug delivery. Nevertheless, the stability of SlpB-LP *in vivo*, and its biodistribution was not investigated.

In this study, we report the biodistribution and stability of SlpB-LP *in vivo*. Stability of SlpB-LP against gut environment was evaluated both *in vitro* and *in vivo*. To quantify the availability of drugs in gut and systemic immune system, concentration of compounds delivered to Peyer’s patches, intestines, and blood were evaluated.

## Methods

### Materials

Lyophilised anionic liposome (Presome PPG-1, DPPC:Chol:DPPG = 1:1:0.2) was obtained from Nippon Fine Chemical (Osaka, Japan); fluorescein isothiocyanate (FITC) was obtained from ICN Biomedicals (California, USA); Cy3 was obtained from BioVision (California, USA); Cy5 was obtained from GE Healthcare (Buckinghamshire, UK); MRS broth was obtained from Becton Dickinson (Maryland, USA); pancreatin was obtained from Amano Enzyme (Nagoya, Japan); 10 K centrifugal filter unit was obtained from Merck Millipore (Cork, Ireland). Other reagents were obtained from Fujifilm Wako Pure Chemicals (Osaka, Japan).

### Extraction of SlpB from *Levilactobacillus brevis* JCM 1059

SlpB was extracted from *Lv. brevis* JCM 1059 and labelled with fluorescent dye when necessary, as described in our prior study [14].

### Preparation of SlpB-Coated Liposome

Lyophilised anionic liposome was dissolved in deionised water by heating at 60°C for 30 min. To prepare fluorescent liposome, either 100 μM FITC or PI was supplemented to each mg of rehydrated liposome; for Cy5 encapsulation, a tube of Cy5 monoreactive dye (fluorophore) was dissolved in 100 μl tris-buffered saline to inhibit conjugation of Cy5 to protein. Then, 6 μL Cy5 was supplemented to each mg rehydrated liposome. Liposome-fluorophore mixture was heated at 60°C for 60 min to facilitate encapsulation. Then, liposome was allowed to cool to room temperature, and excess fluorophore was removed by ultrafiltration with 10 K centrifugal filter unit.

To coat liposome with SlpB, 400 μg/mg [SlpB/LP] was supplemented. The mixture was incubated at 20°C for 120 min with agitation. Then, excess SlpB was removed by centrifugation at 16,000 x*g* for 30 min. To observe SlpB-coating on the surface of anionic liposome, Cy3-SlpB was coated on the surface of FITC-LP, and observed with laser scanning microscope (LSM780, Carl Zeiss, Oberkochen, Germany).

### Zeta potential of SlpB-Coated Liposome

Adsorption of SlpB on surface of liposome will reduce the zeta potential of liposome. Therefore, adsorption capacity of SlpB on liposome can be determined by minimum zeta potential obtained. 10 μg liposome was coated with various concentration of SlpB, and zeta potential was measured with nanoparticle analyser (Horiba, Japan) in a folded capillary cuvette. The relationship between zeta potential and concentration was plotted, and the minimum zeta potential was identified.

### Stability of SlpB-Coated Liposome

Stability of FITC-LP and SlpB-FITC-LP were evaluated in buffer with various pH (pH 2.6, 4.0, 7.0, 9.2), gall solution of various concentrations (0.5%, 1%, 2%, 3%), simulated gut fluid (SGF) and simulated intestinal fluid (SIF) at 37°C for 60 min as described in our prior study [14].

### *In vivo* Stability of SlpB-Coated Liposome

BALB/c mice (female, 12 weeks, 20 – 25 g, Charles River strain) were obtained from Jackson Laboratory, Japan. Animal experiments were approved by the Animal Experiment Committees at Tokyo Institute of Technology (authorization number D2019006-2) and carried out in accordance with the guidelines. Cy5-LP or SlpB-Cy5-LP (200 μL, each containing 82.5 μg liposome) was administered via oral gavage to mice fasted for 3 h (*n* = 4 each group). At 1 h and 3 h administration, the mice were euthanized. Blood, jejunum, ileum, and liver were isolated immediately. Cold PBS (-) were injected into both jejunum and ileum, and the luminal content in both jejunum and ileum were collected. The flow throughs were filled up to equal volume with PBS (-), and the suspensions were allowed to sit on ice for 10 min to allow sedimentation of solid particulate. 1 ml supernatant was sampled and centrifuged at 16,000 x*g* for 30 min within 20 min after sample collection. Then, the supernatants were collected into another tube, and the pellets were resuspended in 1 ml PBS (-). Fluorescence intensity for both supernatant and suspension of pellet was measured, and the fraction of fluorescence intensity of supernatant to fluorescence intensity of pellet was calculated to evaluate the leakage of Cy5 dye from liposome.

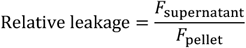

where *F*_supernatant_ indicates fluorescence intensity of supernatant, while *F*_pellet_ indicates fluorescence intensity of pellet. Relative leakage was evaluated instead of relative stability to reduce the uncertainty generated by absorption of liposome.

### Absorption of Liposomes into Intestine and Uptake into Peyer’s Patches

To investigate absorption of liposome into intestine and uptake into Peyer’s patches, 2 fragments of 5 mm intestines from jejunum and ileum without Peyer’s patches, 2 jejunal Peyer’s patches and 2 ileal Peyer’s patches were isolated within 5 min of euthanization. The samples were lysed with 1% polyethylene glycol mono-*p*-isooctylphenyl ether solution with vortex and vigorous shaking, followed by centrifugation at 5,000 x*g* for 5 min. 0.2-volume of supernatants were collected, and the fluorescence intensity was analysed.

### Availability of Fluorophore in Blood

The bloods collected were allowed to sit at room temperature for at least 30 min for clotting. Then, the blood samples were centrifuged at 9,200 x*g* to pellet the blood clot, and the plasma were collected. 10 μL plasma was diluted with 90 μL PBS (-) and the fluorescence intensity was measured. Bioavailability was evaluated by calculating the ratio of fluorescence intensity of fluorophore in blood plasma to initial fluorescence intensity of liposome.

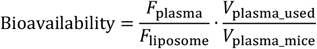

where *F*_plasma_ indicates fluorescence intensity in blood plasma, *F*_liposome_ indicates initial fluorescence intensity of liposome, *V*_plasma_used_ indicates the volume of plasma applied in measurement, *V*_plasma_mice_ indicates the volume of plasma available in mice.

### Availability of Fluorophore in Liver

Livers were collected within 10 min of euthanization of mice. Then, livers were lysed with 1% polyethylene glycol mono-*p*-isooctylphenyl ether solution with vortex and vigorous shaking, followed by centrifugation at 5,000 x*g* for 5 min. 0.2-volume of supernatants were collected, and the fluorescence intensity, and absorbance at 280 nm were measured. Absorbance at 280 nm from 1% polyethylene glycol mono-*p*-isooctylphenyl ether solution was subtracted from absorbance of each sample, and the fluorescence intensity was normalized with absorbance.

## Results

### Preparation of SlpB-Coated Liposome

Coating of SlpB on the surface of liposome was confirmed by observation of FITC-LP and Cy3-SlpB-FITC-LP (Figure 1). Signal of Cy3-SlpB was only detected in Cy3-SlpB-FITC-LP, but not in FITC-LP. Furthermore, Cy3-SlpB was observed at the outer layer of Cy3-SlpB-FITC-LP, suggesting that SlpB was coated on the surface of liposome, but not encapsulated.

**Figure 1.**
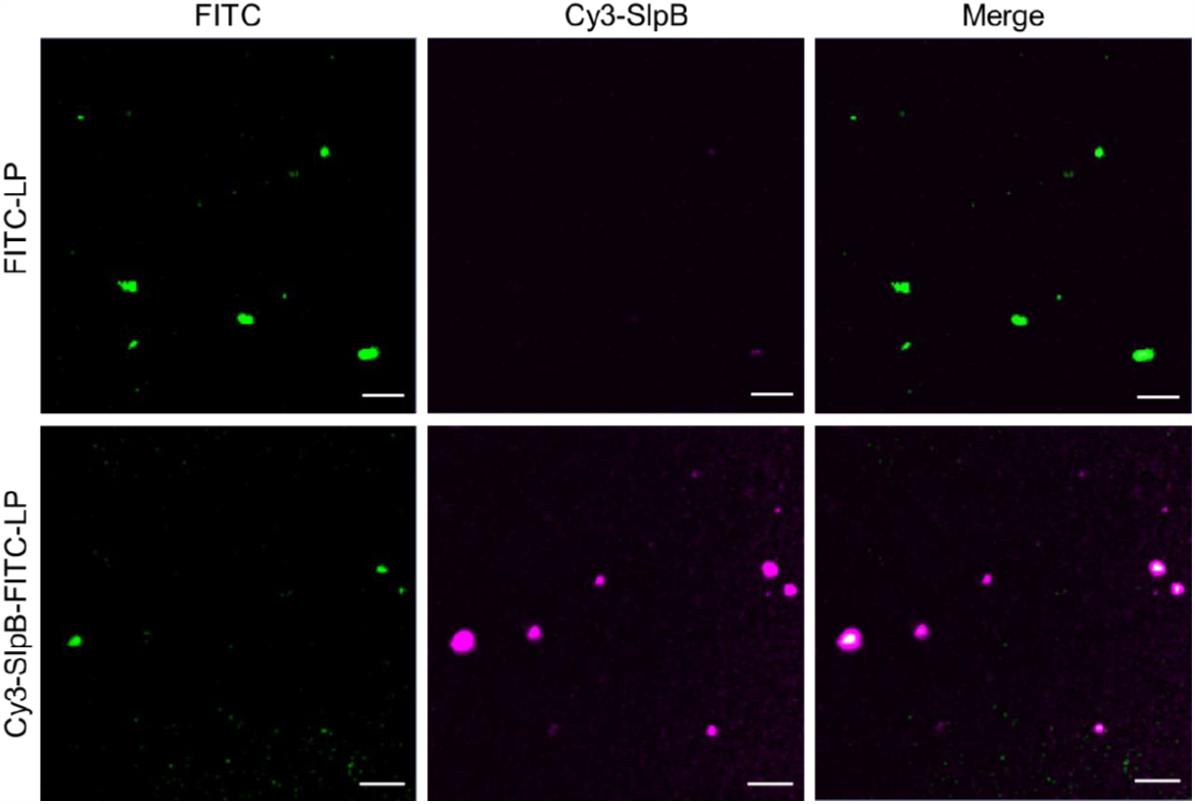
Adsorption of SlpB on liposomes. Fluorescent micrographs of FITC-LP and Cy3-SlpB-FITC-LP. The scale bars represent 5 μm.

### Zeta Potential of SlpB-Coated Liposome

As the composition of liposome used in this study differ from that in our prior study [14], and the size of liposome has been increased from 200 nm to 2 μm, in this study, we have evaluated the adsorption capability of SlpB on liposome through zeta potential measurement, and the stability enhanced by SlpB in relative to bare liposome. Zeta potential of liposome coated with various concentration of SlpB were measured, and an adsorption curve was plotted (Figure 2 (a)). When 1 mg LP was coated with 400 μg SlpB, zeta potential reduced from -19.37 ± 6.19 mV to - 108.27 ± 10.37 mV, which is the minimum zeta potential obtained from liposome coated with 0 – 500 μg/mg [SlpB/LP]. This result suggests that 400 μg SlpB was required to be coated on 1 mg liposome to achieve maximum adsorption capacity.

**Figure 2.**
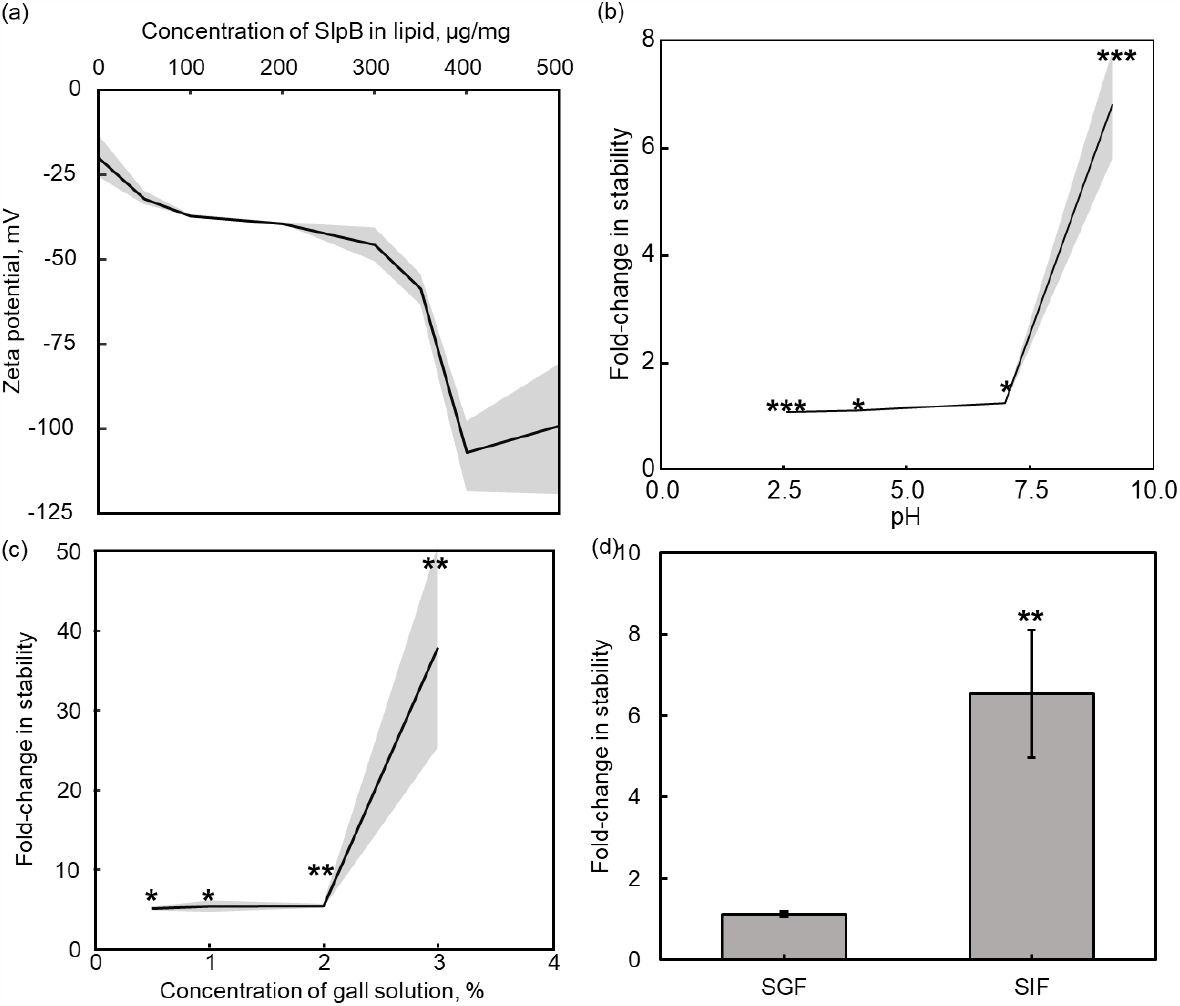
SlpB-coating enhanced the stability of liposome. (a) Zeta potential of liposomes after co-incubation with various concentrations of liposome. Stability of SlpB-LP after incubation in (b) buffers with various pH, (c) gall solution with various concentrations, and (d) simulated gastric fluid and simulated intestinal fluid after incubation at 37°C for 60 min compared to LP. The plots represent mean of data obtained from triplicate samples in independent test and the error bars, and shading regions represent standard deviation of mean. Statistical significance was analysed with Student’s t-test by comparing the data obtained from LP and SlpB-LP. *: *p* < 0.05, **: *p* < 0.01, ***: *p* < 0.001.

### Stability of SlpB-Coated Liposomes

Our previous study suggested SlpB-LP prepared with SlpB at maximum adsorption capacity exhibit highest stability against all condition for monodisperse LP. However, it remained unclear whether this property can be extended to polydisperse LP.

Stability of LP coated with 400 μg/mg [SlpB/LP] was evaluated in various gut mimicking environment. Figure 2 (b – d) suggest that SlpB enhanced stability of polydisperse LP which under various pH, in gall solution with various concentration and SGF, and SIF. At pH 7.00, SlpB-coating has increased the stability of LP by 1.24-fold, from 78.23% FITC retention to 96.92% FITC retention. The effect of stability enhancement was particularly distinctive at pH 9.18 in which the stability of SlpB-LP was enhanced by 6.81-fold from 2.98% to 22.29% (Figure 2 (b)).

For stability in gall solution, SlpB-coating has enhanced stability of LP by 35.41-fold, from 1.95% FITC retention to 68.91% FITC retention (Figure 2 (c)). Although no statistical significance was detected for the enhancement of LP stability by SlpB in SGF, stability of SlpB-LP was increased by 6.54-fold in SIF in relative to LP, from 1.91% FITC retention to 12.47% FITC retention (Figure 2 (d)).

### Stability of SlpB-Coated Liposome *in vivo*

Both drugs and liposomes are subjected to degradation by various enzymatic degradation, mechanical stress, high salinity, and diverse pH environment in gut. To date, stability of liposomes in gut remained unclear due to the difficulty to quantification.

Here, we attempt to evaluate the stability of liposome in gut based on fluorescence intensity of fluorophore. The total fluorescence intensity of liposome conserved if the fluorophore is stable against its environment. In our *in vitro* evaluation, we have shown that the fluorescence intensity of fluorophore is conserved (Figure S1). Thus, by calculating the fraction of fluorescence intensity of supernatant against pellet, we could determine the relative leakage of fluorophore from liposome due to degradation and emulsification of liposome in gut.

As shown in Figure 3, SlpB-coating has significantly reduced the leakage of fluorophore from liposome regardless the length of time after administration. The leakage of fluorophore from SlpB-LP was relatively constant at 1 h and 3 h, where the mean of ratio of fluorescence intensity (RFI) detected were 3.74 and 3.98 respectively compared to 1.61 from control. On the other hand, the mean of RFI detected from intestinal content from mice administered with LP was 5.35- and 6.10-fold higher than SlpB-LP at 1 h and 3 h respectively.

**Figure 3.**
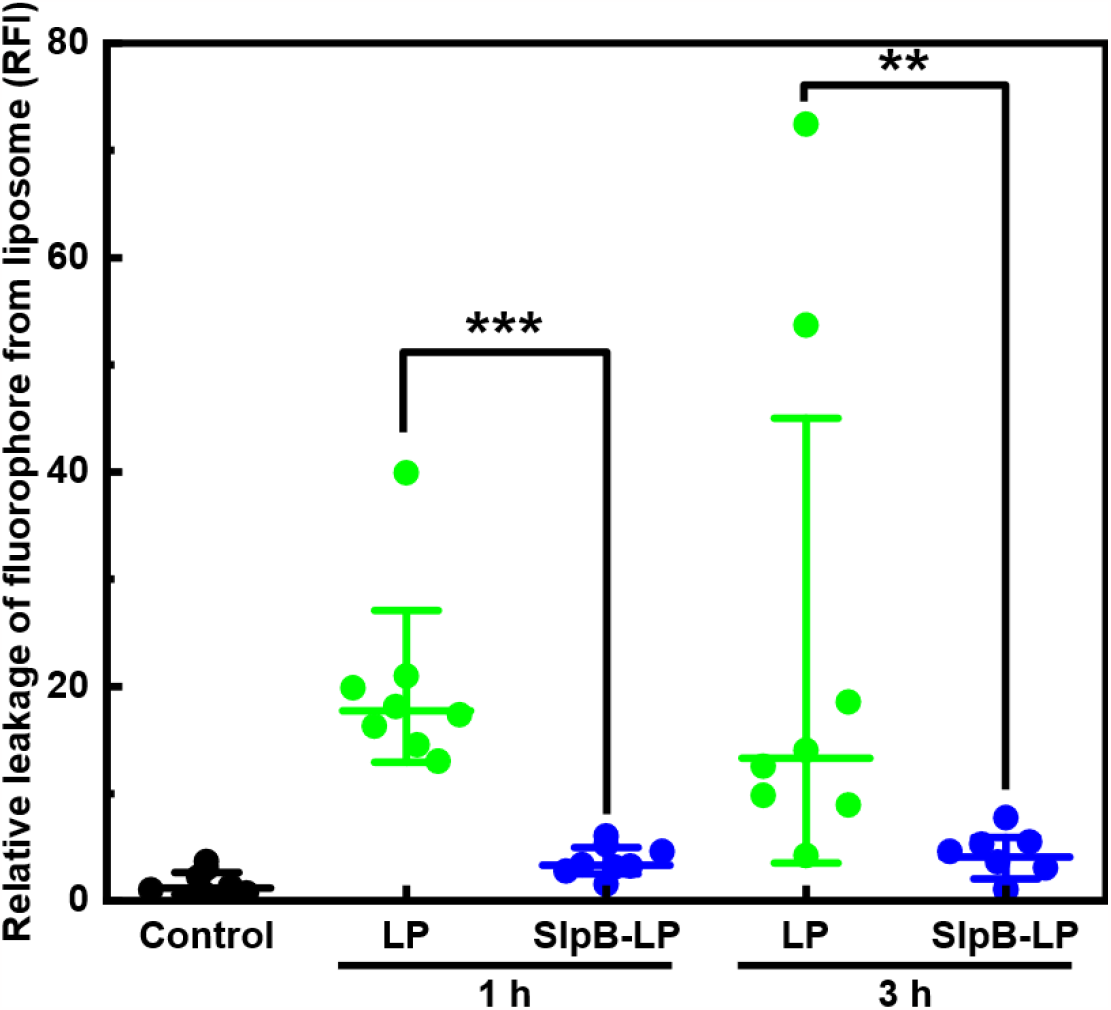
SlpB-coating enhanced the stability of liposome *in vivo*. Ratio of fluorescence intensity obtained from fluorophore leaked into environment to fluorophore remained encapsulated in liposome 1 h and 3 h after oral administration of LP or SlpB-LP to mice. The plots represent median of data obtained from 4 biological replicates and 2 technical replicates and the error bars represents 95% confidence intervals of mean. Statistical significance was analysed with Mann–Whitney U test. ** *p* < 0.01, *** *p* < 0.001.

### Absorption of SlpB-Coated Liposomes into Peyer’s Patches and Intestines

Our previous study has shown that SlpB-coating facilitate transcytosis of liposome into Peyer’s patches through M cells, and enhance specific endocytosis into APCs in Peyer’s patches [14]. However, the specificity of uptake into Peyer’s patches was not clarified. Furthermore, the fate of LP after endocytosis into APCs in Peyer’s patches remained unclear.

By collecting both intestines (sans Peyer’s patches) and Peyer’s patches separately, we were able to evaluate the distribution of LP and SlpB-LP in intestines and Peyer’s patches from the supernatant of tissue lysate. As shown in Figure 4 (a – b), SlpB-LP has been specifically uptake into Peyer’s patches, but not into intestine, while most of the LP were absorbed into intestine. SlpB-coating has increased absorption of LP into Peyer’s patches by 3.60-fold at 1 h, and 2.53-fold at 3 h; while reduced absorption of LP into villi by 8.98-fold at 1 h, and 3.00-fold at 3 h.

**Figure 4.**
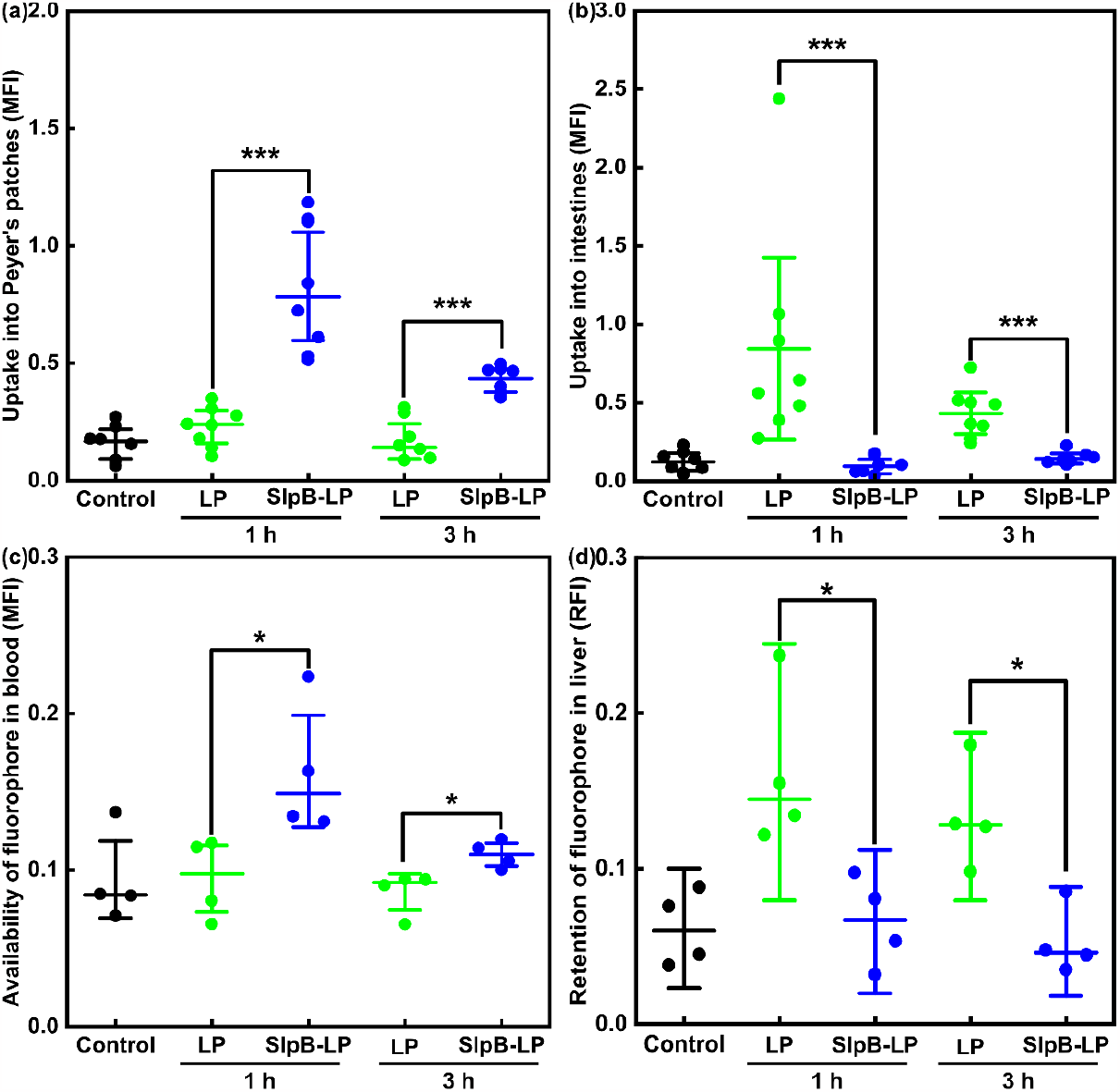
Distribution of fluorophore in mice administered with LP or SlpB-LP. Fluorescence intensity of fluorophore in (a) Peyer’s patches, (b) intestines (sans Peyer’s patches), (c) blood plasma and (d) the ratio of fluorescence intensity to concentration of proteins in liver. The plots represent median of data obtained from 4 biological replicates and 2 technical replicates and the error bars represents 95% confidence intervals of mean for (a) and (b); median of 4 biological replicates and the error bars represents 95% confidence intervals of mean for (c) and (d). Statistical significance was analysed with Mann–Whitney U test. *: *p* < 0.05, ***: *p* < 0.001.

### Availability of SlpB-Coated Liposomes

Orally administered drugs must be transported into lymph or blood vessel to trigger systemic response. To understand whether SlpB-coating could facilitate transportation of LP into systemic circulation, we have measured the fluorescence intensity originated from fluorophore in blood plasma. SlpB-coating has increased the intensity of fluorescence signal from fluorophore in blood plasma by 1.73-fold at 1 h, 1.28-fold at 3 h (Figure 4 (c)). Notably, the fluorescence intensity from blood plasma obtained from mice administered with LP is has no difference from control, which indicates that only a trace amount of fluorophore was available in systemic circulation system. By scaling up the fluorescence intensity from in blood plasma to 0.5 ml, which is the volume of plasma which can be obtained from 20 g mouse, we discovered that availability of fluorophore in blood plasma were 0.017% for mice administered with LP at 1 h, 7.27% and 1.68% for mice administered with SlpB-LP at 1 h and 3 h respectively, while it was below computation limit for mice administered with LP for 3 h. The bioavailability of fluorophore was 427.65-fold higher for mice administered with SlpB-LP compared to LP 1 h after administration.

Then, we have measured the fluorescence intensity in supernatant of lysate of liver (Figure 4 (d)). Fluorescence intensity was reduced by 2.46-fold at 1 h, and 2.51-fold at 3 h when SlpB was coated on LP.

Interestingly, we have found that uptake into Peyer’s patches was correlated to the availability of fluorophore in blood plasma (Figure 5). The coefficient of correlation between the 2 parameters was 0.70. On the other hand, the absorption into intestine was correlated to distribution of fluorophore in liver, with coefficient of correlation of 0.68 (Figure S2).

**Figure 5.**
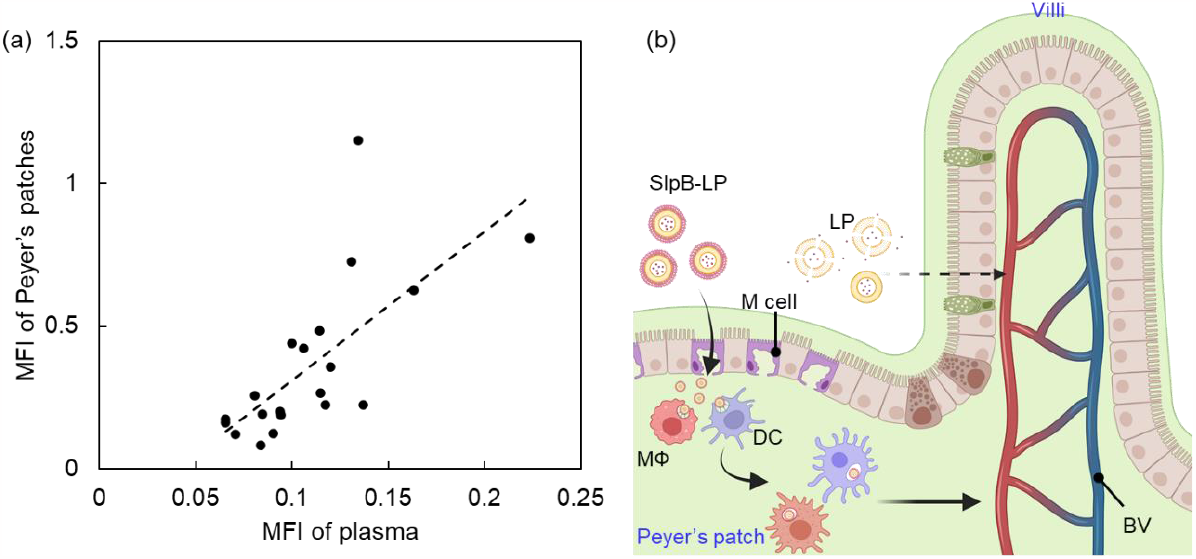
Relationship of concentration of fluorophores in Peyer’s patches and blood plasma. (a) Scatter plot of fluorescence intensity of fluorophore in Peyer’s patches against fluorescence intensity of fluorophore in blood plasma. Linear regression was performed, and a linear curve was plotted. The coefficient of correlation is 0.70. (b) Illustration of transportation of SlpB-LP in gut. SlpB-LP was transported by APCs in Peyer’s patches into blood vessels. BV: blood vessel, DC: dendritic cell, LP: liposome, MΦ: macrophage, SlpB-LP: SlpB-coated liposome.

## Discussion

Prior studies have reported that the adsorption capacity of SlpB on cationic LP was 200 μg/mg [SlpB/LP] [11]. The adsorption capacity of SlpB on the surface of anionic LP remained uncertain. By using anionic liposome with different composition compared to our prior study, which the DPPC:Chol:DPPG ratio was 1:1:0.2 (this study) compared to 0.75:1:0.75 (previous study) [14], we have found that the maximum adsorption capacity of SlpB was 400 μg/mg [SlpB/LP] on anionic liposome (Figure 2 (a)), regardless its composition, particle size, and initial zeta potential. The adsorption capacity of SlpB on anionic liposome is 2-times higher than cationic liposome, probably due to the formation of denser crystalline array on the surface of anionic liposome, which the charge is similar to native cell wall of *Lv. brevis*. Further investigation of the formation of crystalline array and its geometry has to be conducted.

Slp binds to and rigidify the lipid membrane of liposome. This interaction of Slp with lipid enhances the stability of liposome by increasing the rigidity of lipid membrane [15]. Furthermore, formation of crystalline array on LP surface could protect LP from enzymatic hydrolysis. In this study, we have shown that SlpB could enhance the stability of liposome regardless of size and polydispersity (Figure 2 (b – d)). This property is useful in delivery of biological lipid vesicles, e.g., extracellular vesicles, which the size are indefinite and sometimes exist in μm scale. The protective effect of SlpB on liposome is particularly distinctive under lipid degrading conditions, i.e., in 3% gall solution which can emulsify lipid and SIF which contains lipase and protease (in pancreatin).

Notably, SlpB has enhanced stability of liposome in gut environment (Figure 3). In gut, mechanical stress, enzymatic degradation, e.g., pepsin hydrolysis, protease hydrolysis, lipase hydrolysis, constant changes in pH, and emulsification could result in degradation of liposome. SlpB has formed a protective layer on the surface of liposome, while rigidifying lipid membrane, which have increased the stability of LP in gut 5.35- and 6.10-fold at 1 h and 3 h after oral administration.

SlpB could target M cells and APCs in Peyer’s patches [14]. The targeting capability has enhanced uptake of liposome into Peyer’s patches (Figure 4 (a)). With Slp from *L. helveticus*, previous study has suggested that Slp from *L. acidophilus* group binds to and release the content of liposome to mucosal layer of gastrointestinal tract [16]. Although this approach is a fascinating in vaccine delivery, it does not enhance drug delivery to APCs in Peyer’s patches, and weak in inducing systemic immune response. In our study, we discovered that the fluorescence intensity in intestines for mice administered with SlpB-LP did not increase in comparison with control, which suggested that SlpB-LP did not release its content in mucosal layer. This result might be due to the existence of both extracellular matrix binding site [17], sugar-binding site and cell wall binding domain at N-terminal for Slp from *Lv. brevis-*group lactobacilli. As the binding sites are covered with crystal forming C-terminal, access of these binding sites to mucosal layer might be blocked, thus reducing the interaction to mucosal layer. This result further confirm that the orientation of SlpB on LP is similar to those on bacterial surface. On the other hand, the mucin binding site of SlpA from *L. acidophilus* group bacteria was found to be localised at crystalline array forming N-terminal, which is in opposite direction to cell wall binding domain at C-terminal. This property results in unspecific binding of Slp to extracellular matrices, e.g., mucosal layer, which result in release of drugs in mucosal layer.

For mice administered with LP, 2 reasons contributed to the increases in fluorescence intensity in intestine. Without SlpB-coating, (1) unspecific uptake of LP occurred and (2) higher fraction of LP degradation resulted in higher fraction of leaked fluorophore (Figure 3). As villi has a larger surface layer to volume ratio, higher concentration of LP and leaked fluorophore were absorbed into intestines through intestinal villi compared to Peyer’s patches (Figure 4 (b)).

Interestingly, delivery through Peyer’s patches has increased the concentration of fluorophore in blood plasma (Figure 4 (c)). Concentration of fluorophores in Peyer’s patches is correlated to concentration in blood plasma (Figure 5 (a)), which suggest that delivery through Peyer’s patches might be an important route for drugs delivery to systemic circulation (Figure 5 (b)). The increase in bioavailability of compound might resulted from the capability to bypass first pass elimination in liver (Figure 4 (d)) via Peyer’s patches targeting. On the other hand, absorption through intestine resulted in metabolism in intestinal wall, transportation of fluorophore to liver through portal vein, and metabolism in liver, which has been shown by high concentration of fluorophore in both intestine and liver for mice administered with LP.

In conclusion, we have demonstrated that SlpB-coated liposome is a promising active drug delivery system to target APCs in Peyer’s patches and induced both localised and systemic immune responses.

## Author Contributions

NY and ZLT contributed to all processes of this study.

## Ethics Approval

Animal experiments were approved by the Animal Experiment Committees at Tokyo Institute of Technology (authorization number D2019006-2) and carried out in accordance with the guidelines.

## Reference

1. Corr SC, Gahan Ccgm, Hill C. M-cells: origin, morphology and role in mucosal immunity and microbial pathogenesis. FEMS Immunol Med Microbiol. 2008;52:2–12.

2. Jung C, Hugot J-P, Barreau F. Peyer’s Patches: The Immune Sensors of the Intestine. Int J Inflamm. 2010;2010:1–12.

3. Pond SM, Tozer TN. First-Pass Elimination: Basic Concepts and Clinical Consequences. Clin Pharmacokinet. 1984;9:1–25.

4. Sosnik A, Augustine R. Challenges in oral drug delivery of antiretrovirals and the innovative strategies to overcome them. Adv Drug Deliv Rev. 2016;103:105–20.

5. Chen Y, Wu J, Wang J, Zhang W, Xu B, Xu X, et al. Targeted delivery of antigen to intestinal dendritic cells induces oral tolerance and prevents autoimmune diabetes in NOD mice. Diabetologia. 2018;61:1384–96.

6. Yang J, Sun H, Song C. Preparation, characterization and in vivo evaluation of pH-sensitive oral insulin-loaded poly(lactic-co-glycolicacid) nanoparticles. Diabetes Obes Metab. 2012;14:358–64.

7. Song JG, Lee SH, Han H-K. Development of an M cell targeted nanocomposite system for effective oral protein delivery: preparation, in vitro and in vivo characterization. J Nanobiotechnology. 2021;19:15.

8. Lin S, Mukherjee S, Li J, Hou W, Pan C, Liu J. Mucosal immunity–mediated modulation of the gut microbiome by oral delivery of probiotics into Peyer’s patches. Sci Adv. 2021;7:eabf0677.

9. Yano A, Takahashi K, Mori Y, Watanabe S, Hanamura Y, Sugiyama T, et al. Peyer’s Patches as a Portal for DNA Delivery by Lactococcus lactis in Vivo. Biol Pharm Bull. 2018;41:190–7.

10. Rios D, Wood MB, Li J, Chassaing B, Gewirtz AT, Williams IR. Antigen sampling by intestinal M cells is the principal pathway initiating mucosal IgA production to commensal enteric bacteria. Mucosal Immunol. 2016;9:907–16.

11. Hollmann A, Delfederico L, Glikmann G, De Antoni G, Semorile L, Disalvo EA. Characterization of liposomes coated with S-layer proteins from lactobacilli. Biochim Biophys Acta BBA - Biomembr. 2007;1768:393–400.

12. Konstantinov SR, Smidt H, de Vos WM, Bruijns SCM, Singh SK, Valence F, et al. S layer protein A of Lactobacillus acidophilus NCFM regulates immature dendritic cell and T cell functions. Proc Natl Acad Sci. 2008;105:19474–9.

13. Yanagihara S, Kanaya T, Fukuda S, Nakato G, Hanazato M, Wu X-R, et al. Uromodulin– SlpA binding dictates Lactobacillus acidophilus uptake by intestinal epithelial M cells. Int Immunol. 2017;29:357–63.

14. Tan ZL, Kitamoto Y, Miyanaga K, Yamamoto N. Levilactobacillus brevis surface layer protein B promotes liposome targeting to antigen-presenting cells in Peyer’s patches. Int J Pharm. 2022;622:121896.

15. Hollmann A, Delfederico L, De Antoni G, Semorile L, Disalvo EA. Interaction of bacterial surface layer proteins with lipid membranes: Synergysm between surface charge density and chain packing. Colloids Surf B Biointerfaces. 2010;79:191–7.

16. Wang W, Shao A, Feng S, Ding M, Luo G. Physicochemical characterization and gastrointestinal adhesion of S-layer proteins-coating liposomes. Int J Pharm. 2017;529:227–37.

17. Hynönen U, Westerlund-Wikström B, Palva A, Korhonen TK. Identification by Flagellum Display of an Epithelial Cell- and Fibronectin-Binding Function in the SlpA Surface Protein of Lactobacillus brevis. J Bacteriol. 2002;184:3360–7.

