## Supplementary material for "*In vivo* Stability and Biodistribution of Liposome Coated with SlpB from *Levilactobacillus brevis*": Figure S1; Figure S2

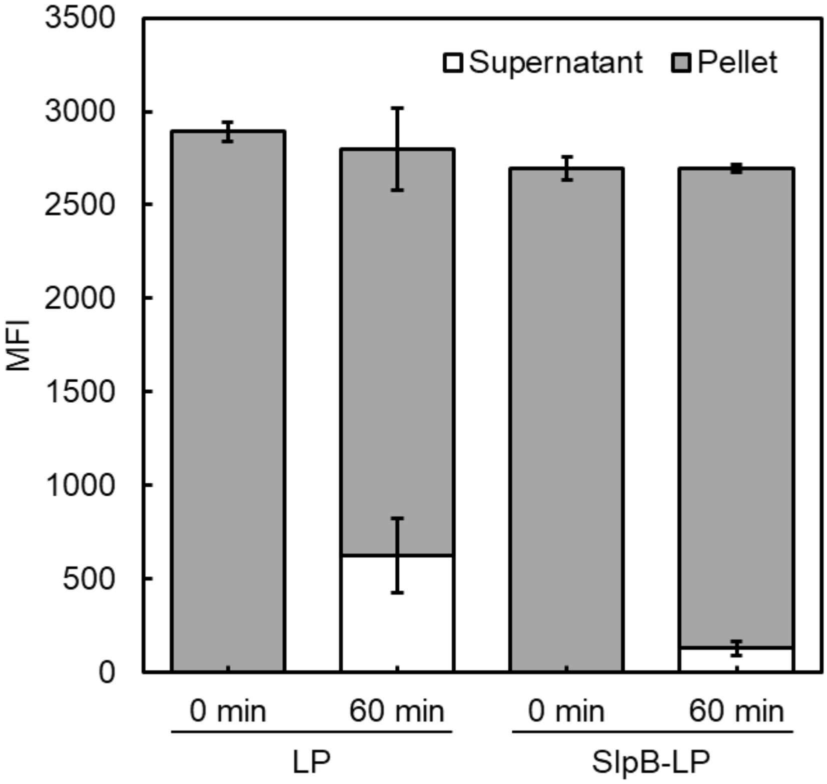


**Figure S1 Conservation of fluorescent dye in whole sample.** After incubation in buffer with pH 7 for 60 min at 37°C, the liposomes were centrifuged at 16,000 xg for 30 min. Then, the supernatant was collected, and the pellet was resuspended with equal volume of PBS (-). Fluorescence intensity of sample before incubation, supernatant and pellet after incubation were measured. The sum of fluorescence intensity of supernatant and pellet after incubation (60 min) were compared with fluorescence intensity of sample before incubation (0 min). The plots represent data obtained from triplicate samples in independent test and the error bars represent standard deviation of mean.

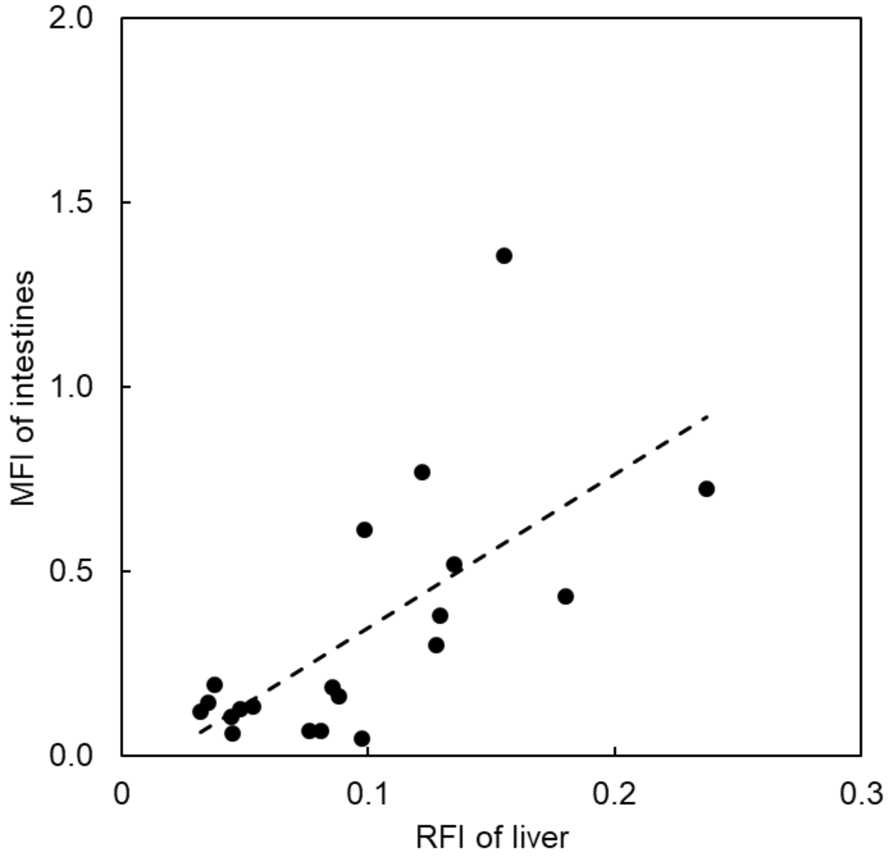


**Figure S2 Relationship of concentration of fluorophores in intestines and liver.** Scatter plot of fluorescence intensity of fluorophore in intestines against fluorescence intensity of fluorophore in liver. Linear regression was performed, and a linear curve was plotted. The coefficient of correlation is 0.68.
